# Comparative genomics and the salivary transcriptome of the redbanded stink bug shed light on its high damage potential to soybean

**DOI:** 10.1101/2023.12.15.571890

**Authors:** Hunter K. Walt, Jonas G. King, Tyler B. Towles, Seung-Joon Ahn, Federico G. Hoffmann

**Affiliations:** Department of Biochemistry, Molecular Biology, Entomology, and Plant Pathology, Mississippi State University, Mississippi State, MS 39762, USA; Macon Ridge Research Station, Louisiana State University, Winnsboro, LA 71295, USA; Institute for Genomics, Biocomputing and Biotechnology, Mississippi State University, Mississippi State, MS 39762, USA

**Author notes:** Author for Correspondence: Federico G. Hoffmann, Department of Biochemistry, Molecular Biology, Entomology, and Plant Pathology, Mississippi State University, Mississippi State, MS 39762, USA.

**Keywords:** *Piezodorus guildinii*, gene family evolution, Hemiptera, gene loss, gene duplication, differential retention

## Abstract

The redbanded stink bug, *Piezodorus guildinii* (Westwood) (Hemiptera: Pentatomidae), is a significant soybean pest in the Americas, inflicting more physical damage on soybean than other native stink bugs. Studies suggest that its heightened impact is attributed to the aggressive digestive properties of its saliva. Despite its agricultural importance, the factors driving its greater ability to degrade plant tissues have remained unexplored in a genomic evolutionary context. In this study, we hypothesized that lineage-specific gene family expansions have increased the dosage of digestive genes expressed in the salivary glands. To investigate this, we annotated a previously published genome assembly of the redbanded stink bug and performed a comparative genomic analysis on 11 hemipteran species and reconstructed patterns of gene duplication, gain, and loss in the redbanded stink bug. We also performed RNA-seq on the redbanded stink bug’s salivary tissues, along with the rest of the body without salivary glands. We identified hundreds of differentially expressed salivary genes, including a subset lost in other stink bug lineages but retained and expressed in the redbanded stink bug’s salivary glands. These genes were significantly enriched with protein families involved in proteolysis, potentially explaining the redbanded stink bug’s heightened damage to soybeans. Contrary to our hypothesis, we found no support for an increased dosage of digestive genes in the salivary glands of the redbanded stink bug. Nonetheless, these results provide insight into the evolution of this important crop pest, establishing a link between its genomic history and its agriculturally important physiology.

**SIGNIFICANCE STATEMENT:** The redbanded stink bug, an important soybean pest in the Americas, inflicts greater damage to soybean due to higher salivary digestion. We employed comparative genomics and analyses of the salivary transcriptome to explore this and found that the differential retention of ancestral salivary genes may explain this phenomenon better than gene gains or duplications in the redbanded stink bug genome. We identify a distinct set of genes in the salivary gland that were differentially retained and expressed in the redbanded stink bug lineage, demonstrating enrichment in proteolytic function. This discovery offers a potential explanation for the redbanded stink bug’s elevated damage to soybean crops.

## INTRODUCTION

The redbanded stink bug, *Piezodorus guildinii* (Westwood) (Hemiptera: Pentatomidae), is an important soybean (*Glycine max*, L.) pest in the Americas with a distinct ability to digest plant tissues. It was first described in 1837 on the island St. Vincent in the Caribbean, and although its geographic origin remains unclear, studies have suggested that it originated in the Caribbean Basin and reached Brazil about a million years ago (Moraes et al. 2023). Then, more recently, the redbanded stinkbug expanded its range and population size in conjunction with increases in soybean production across South America (Moraes et al. 2023; Zucchi et al. 2019; Bundy et al. 2018). Now, it ranges from Argentina through Central America, Mexico, and the southern United States from Texas to South Carolina (Sosa-Gómez et al. 2020). It has been consistently designated as an economically significant pest in Brazil, Argentina, and Uruguay since the 1970s (more recently in the United States), and it is highly capable of expanding its range (Chen et al. 2023; Bundy et al. 2018). Interestingly, other species of the genus *Piezodorus* are pests of legume crops in Europe, Asia, Africa, and Australia, but the redbanded stink bug is the only species that occurs in the New World, with its closest relatives, *P. oceanicus* and *P. hybneri*, occurring in Australia, Asia and Africa respectively (Bundy et al. 2018) (**Supplementary Figure 1**).

The redbanded stink bug mainly feeds on the pods of legumes, causing significantly more damage to soybean seeds than other stink bug species (Depieri & Panizzi 2011; Parker 2012; Husch et al. 2014; Tuelher et al. 2016; Bundy et al. 2018). When the insect feeds on plants, its salivary glands release a concoction of proteins that digest plant tissues, detoxify harmful plant compounds, and mitigate plant defense responses. Consequently, the study of salivary tissues becomes crucial in the context of crop pests (Sharma et al. 2014; Marshall et al. 2023). Previous studies have shown that when the redbanded stink bug feeds on soybeans, there is a higher level of chemical dissolution compared to other stink bugs, suggesting an increased digestive component in their saliva (Corrêa-Ferreira & De Azevedo 2002; Depieri & Panizzi 2011; Husch et al. 2014). Additionally, it has been noted that the redbanded stink bug exhibits a preference for plants in the Fabaceae family (legumes), and is rarely observed on crops other than soybean, unlike other relevant stink bug pests that feed on diverse crops such as cotton and corn (Temple et al. 2013a). Furthermore, resistance to insecticides has been documented in the redbanded stink bug (Baur et al. 2010). One study has suggested that the redbanded stink bug is less susceptible to pyrethroids and organophosphates than the southern green stink bug (Temple et al. 2013b).

The genetic underpinnings of the greater damage inflicted to soybean by the redbanded stink bug are unknown, and we hypothesize that changes in the redbanded stink bug genome may account for this distinct phenotype. Specifically, we predict that (1) the repertoire of digestion-related genes would have expanded in the redbanded stink bug genome, and that (2) these genes would exhibit high expression levels in the salivary glands. Gene duplication (i.e., when a gene has an increase in copy number) and subsequent mutation accumulation can lead to new or more specialized protein functions (Lynch & Conery 2000). These novel functions can be retained if they confer a fitness benefit that makes an organism better adapted to its environment (Pearce et al. 2017). In addition to providing fuel for evolutionary novelties, an increase in gene dosage can lead to higher expression levels of a gene family, thus increasing the likelihood of translation of those proteins (Ragipani et al. 2022). Along with this, changes in gene expression at particular loci can facilitate an organism’s adaptation to its surroundings (López-Maury et al. 2008; Nourmohammad et al. 2017). To test our predictions, we compared the genomes of multiple hemipteran taxa, including a chromosome-level genome for the redbanded stink bug (Saha et al. 2022). Additionally, we generated transcriptomic data from field-collected redbanded stink bugs to explore gene expression patterns in the salivary gland components, namely the principal salivary gland (PSG), the accessory salivary gland (ASG), and the accessory salivary ducts (ASD) (**Figure 1**). We used these data to investigate patterns of gene gain, duplication, and loss in the redbanded stink bug lineage and ultimately identify the genes that could account for its heightened damage potential to soybeans.

**Figure 1:**
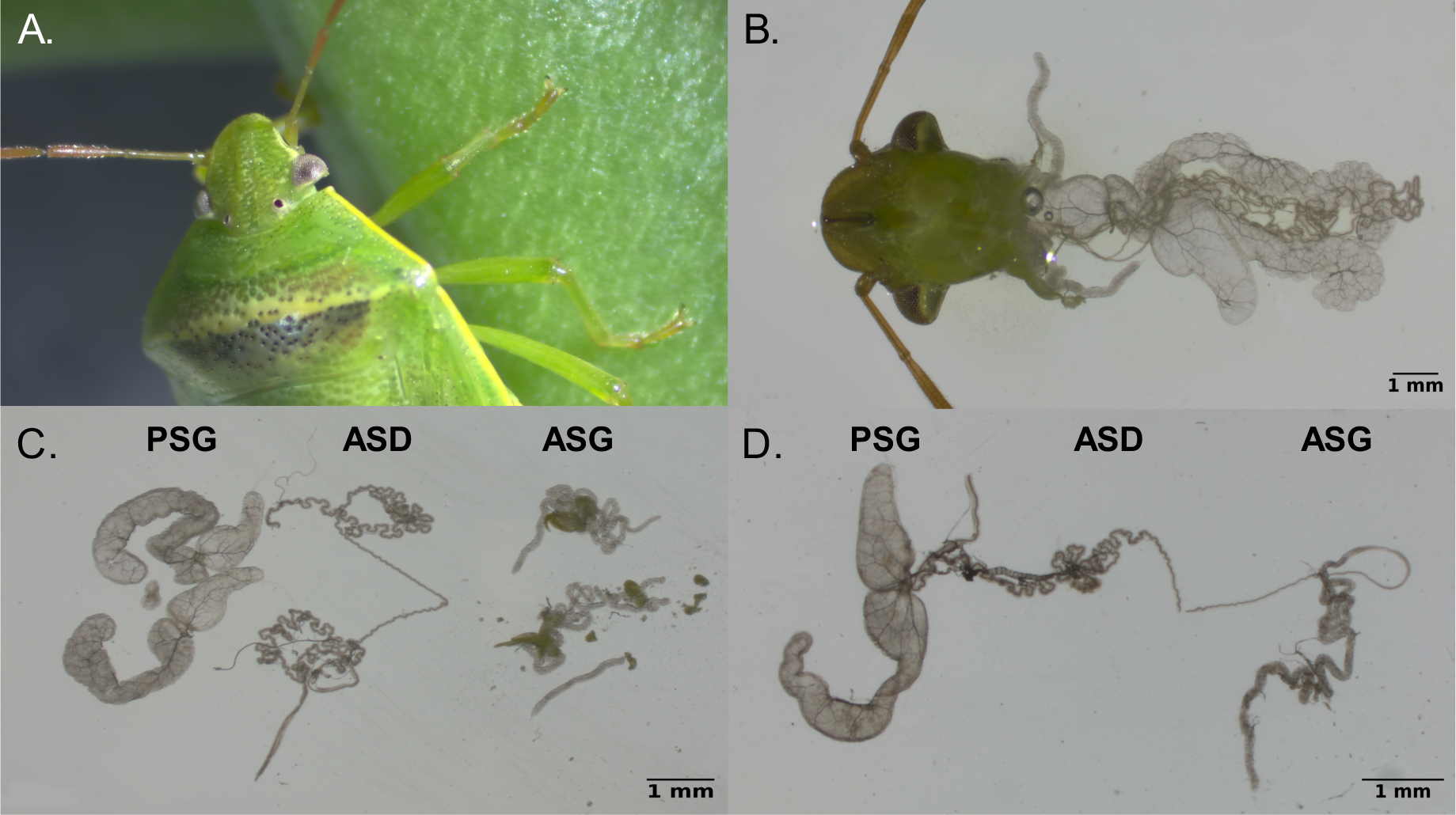
Photographs of the redbanded stink bug and its salivary tissues. **A.)** A photograph of a redbanded stink bug feeding on a green bean. **B.)** Ventral view of a redbanded stink bug head with its salivary tissues displayed. **C.)** Fully dissected salivary tissues labeled principal salivary gland, (PSG), accessory salivary duct (ASD), and accessory salivary gland (ASG). **D.)** Mostly intact salivary tissues with their corresponding labels. The ASG and PSG are connected via the ASD.

## RESULTS

### Genome Annotation

We identified 62.09% of the redbanded stink bug genome (Genbank: GCA_023052935.1) as repetitive elements. These regions were masked, and we annotated the masked genome using the BRAKER2 pipeline. We compared all predicted proteomes from RNA-seq data and protein data before and after TSEBRA transcript selection (see MATERIALS AND METHODS), and we found the highest BUSCO score with the genome annotation using RNA-seq hints and Augustus gene prediction. We ran the supplementary BRAKER script, selectSupportedSubsets.py, to filter the predicted genes by any external support. This resulted in 13,563 annotated genes with support, of which 12,641 (93.2%) have functional annotations. We ran BUSCO analysis in the protein mode using the hemiptera_odb10 database and found that 2,144 universal single-copy orthologs out of 2,510 (85.4%) were present in the final protein dataset (**Supplementary Table 1**).

### Comparative Genomics

We used the 11 hemipteran genomes that met our thresholds for our comparative genomics analysis (**Supplementary Table 1**). A total of 196,495 genes were assigned to 17,545 orthogroups by the OrthoFinder algorithm. We found 1,339 single-copy orthologs shared between the 11 species, and we built a species tree based on a concatenated alignment of the single-copy orthologs (**Figure 2**).

**Figure 2:**
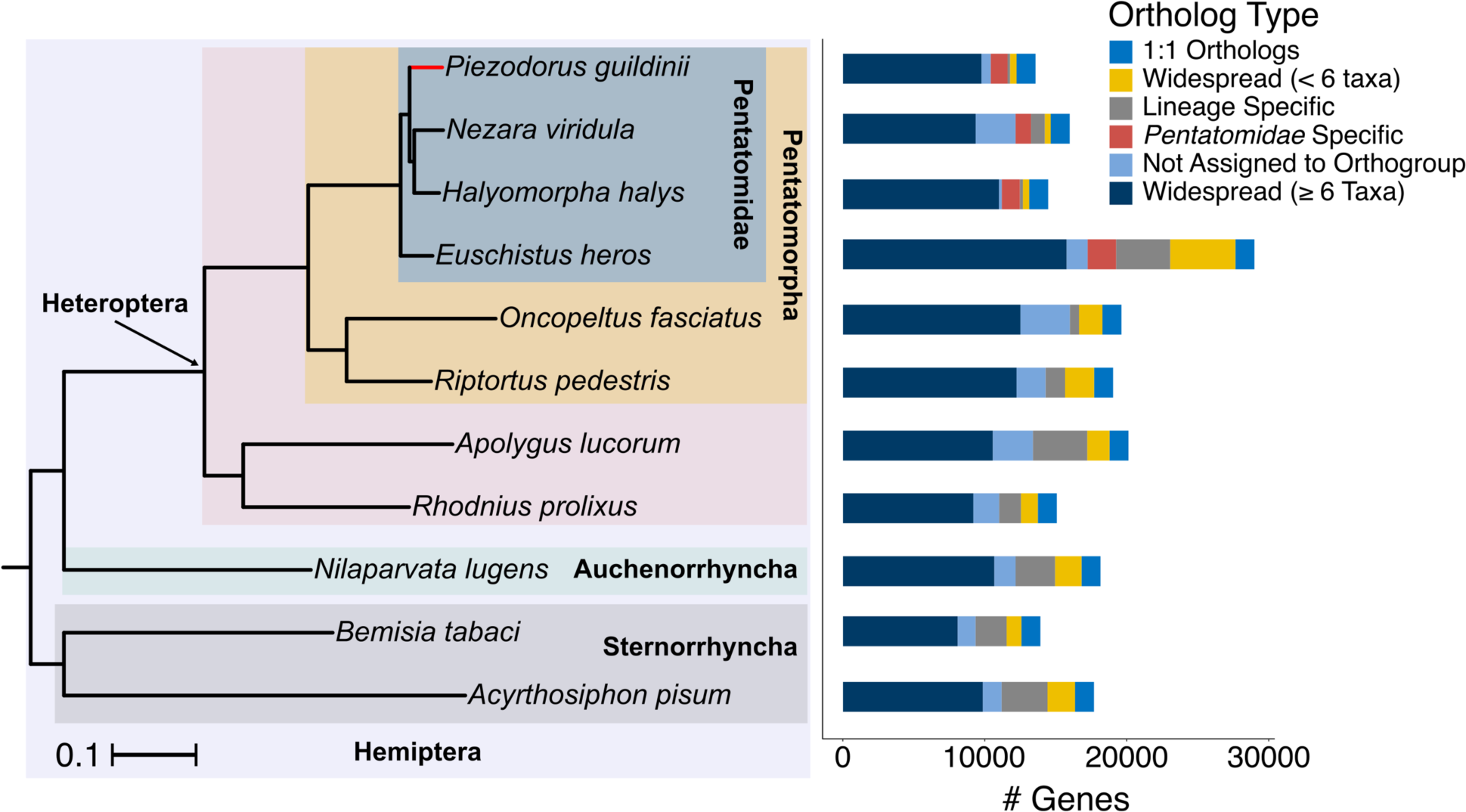
Species tree of the hemipteran taxa used in this study and comparative genomics overview. The species tree is based on a concatenated alignment of 1,339 single-copy orthologs identified by OrthoFinder. All nodes had 100% support. The comparative genomics overview bars correspond to the branches on the tree. The overview is based on gene classification from OrthoFinder, where we further categorized orthologs based on the number of taxa that they are present in.

Because studies have shown that the redbanded stink bug causes more damage to soybean than other stink bugs, we used our comparative genomics analysis to identify changes in gene families could be important to this physiology. Using OMA standalone and pyHam, we identified 618 duplicated genes, 2,451 gained genes, and 4,493 lost genes in the redbanded stink bug lineage. Of note, when considering “gained” genes assigned by OMA and pyHam, two things should be considered. (1) OMA is relatively stringent in its orthogroup assignment, and thus “gained” genes may be highly diverged duplicated genes, not necessarily orphan genes (Altenhoff et al. 2016). (2) The number of gained genes in extant taxa may be inflated by errors in genome assembly (Zajac et al. 2021). We conducted KEGG orthology term enrichment analysis on duplicated, gained, and lost genes identified by pyHam from the redbanded stink bug lineage using its annotated genome as the background. We found 35 and 3 significantly enriched KEGG orthology terms in the duplicated and gained genes, respectively (**Figure 3A**). We found significant enrichment of gene families involved in detoxification and protein digestion in the set of duplicated genes (**Figure 3A**). The three significantly enriched KEGG orthology terms in the set of gained genes were, K13872, K13867, and K10408, which correspond to solute carrier family 7 protein members 6, 7 and dynein axonemal heavy chain, respectively, with adjusted p-values of 1.78e-08, 1.78e-08 and 0.021. To identify enriched KEGG orthology terms in the genes lost in the redbanded stink bug lineage, we used pyHam to infer the genome of the direct ancestor of the redbanded stink bug. The inferred genome had an estimated 15,208 genes, of which 14,253 (93.7%) were functionally annotated by eggNOG. We used pyHam to obtain the extant orthologs of the genes lost in the redbanded stink bug lineage and used these genes to test for enrichment of KEGG orthology terms. We detected 12 significantly enriched KEGG orthology terms lost in the redbanded stink bug lineage (**Figure 3B**).

**Figure 3:**
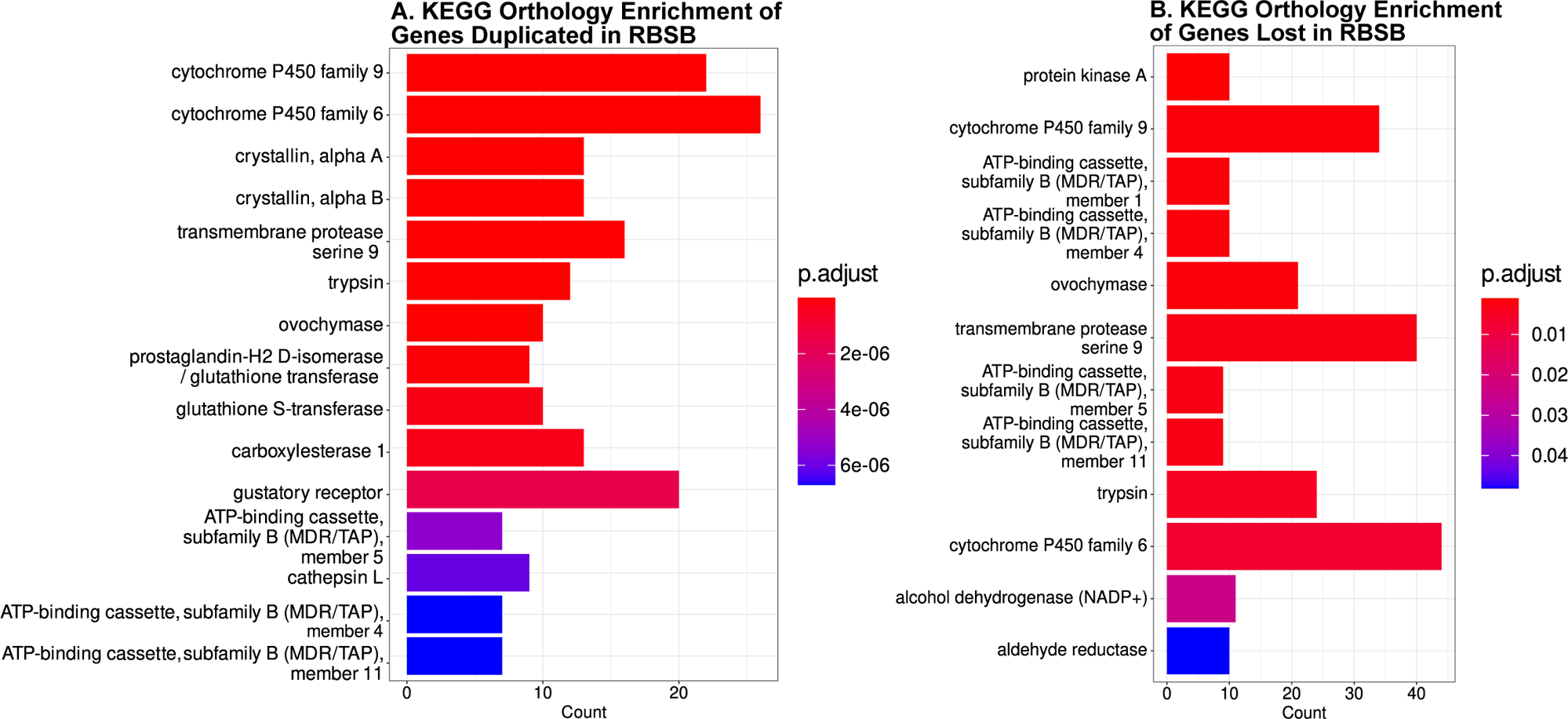
Enrichment of KEGG orthology terms in the genes that were duplicated and lost in the redbanded stink bug (RBSB) lineage. **A.)** Top 10 KEGG orthology terms significantly enriched in duplicated genes in the RBSB lineage. The entire annotated genome was used as the background set of genes. We found functional ortholog enrichment of genes involved in detoxification, such as cytochrome P450s, glutathione S-transferases, carboxylesterases, and ABC transporters, and protein degradation such as trypsins, ovochymases, and cathepsins. **B.)** KEGG orthology terms significantly enriched in genes lost in the redbanded stink bug lineage. The annotated genome of the redbanded stink bug’s direct ancestor (inferred from pyHam) was used as the background set of genes, and the genes that were lost in the redbanded stink bug lineage but retained in the brown marmorated stink bug or southern green stink bug were used as the query. All significantly enriched terms are shown (*n* = 12). Interestingly, we found similar functional enrichment in duplicated and lost genes. There were only three gene families that were overrepresented in the set of gained genes, which are stated in the results section.

To test if the redbanded stink bug lineage had a higher rate of gene turnover than the rest of the taxa in this study, we used CAFE5 to determine separate gene turnover rates (λ) for 1.) the redbanded stink bug lineage (λ_rbsb_), 2.) the rest of the pentatomids (λ_p_), and 3.) the rest of the Hemiptera (λ_h_). We found that the redbanded stink bug lineage and the rest of the pentatomids had similar turnover rates, which was slightly lower in the redbanded stink bug than in the rest of the pentatomids (λ_rbsb_ = 0.0022 and λ_p_ = 0.0026, respectively). The rest of the Hemiptera had nearly a three-fold lower turnover rate (λ_h_ = 0.0009) relative to both λ_rbsb_ and λ_p_. The models included 4 discrete gamma categories and an estimated error value of 0.05587. The negative log-likelihood of the final model was 113,818.

### Differential Expression

We first confirmed that sex does not account for many significant differences between our tissues (**Supplementary Figure 2**). There were only 40 genes that were differentially expressed between males and females. Of these, 25 were upregulated in males and 15 were upregulated in females with many being expected (e.g., *doublesex* and vitellogenin protein genes upregulated in females). Thus, we did not account for sex in the rest of our analyses.

We found 896, 366, and 705 genes significantly upregulated in the PSG, ASG, and ASD tissues, respectively, versus 2,900, 1,022, and 2,889 genes downregulated (**Figure 4A-C**). It is noteworthy that there is minimal overlap among differentially expressed genes across these salivary tissues (**Figure 4D**). We assessed GO term enrichment of the differentially expressed genes from each salivary tissue and found 21 molecular function GO terms enriched in the PSG, 25 in the ASG, and 13 in the ASD (**Tables 1-3**). In the PSG, there was significant enrichment of functions related to the degradation of proteins such as endopeptidases, exopeptidases, and serine hydrolases (**Table 1**). In the ASG, we found significant enrichment of functions including protein and nucleotide binding, transporter activity, and GTPase activity (**Table 2**). In the ASD, we found enriched functions involving proton transport, ATP hydrolysis, phosphotransferase activity, and lyase activity (**Table 3**). These results suggest that most of the digestive enzymes are produced in the PSG.

**Figure 4:**
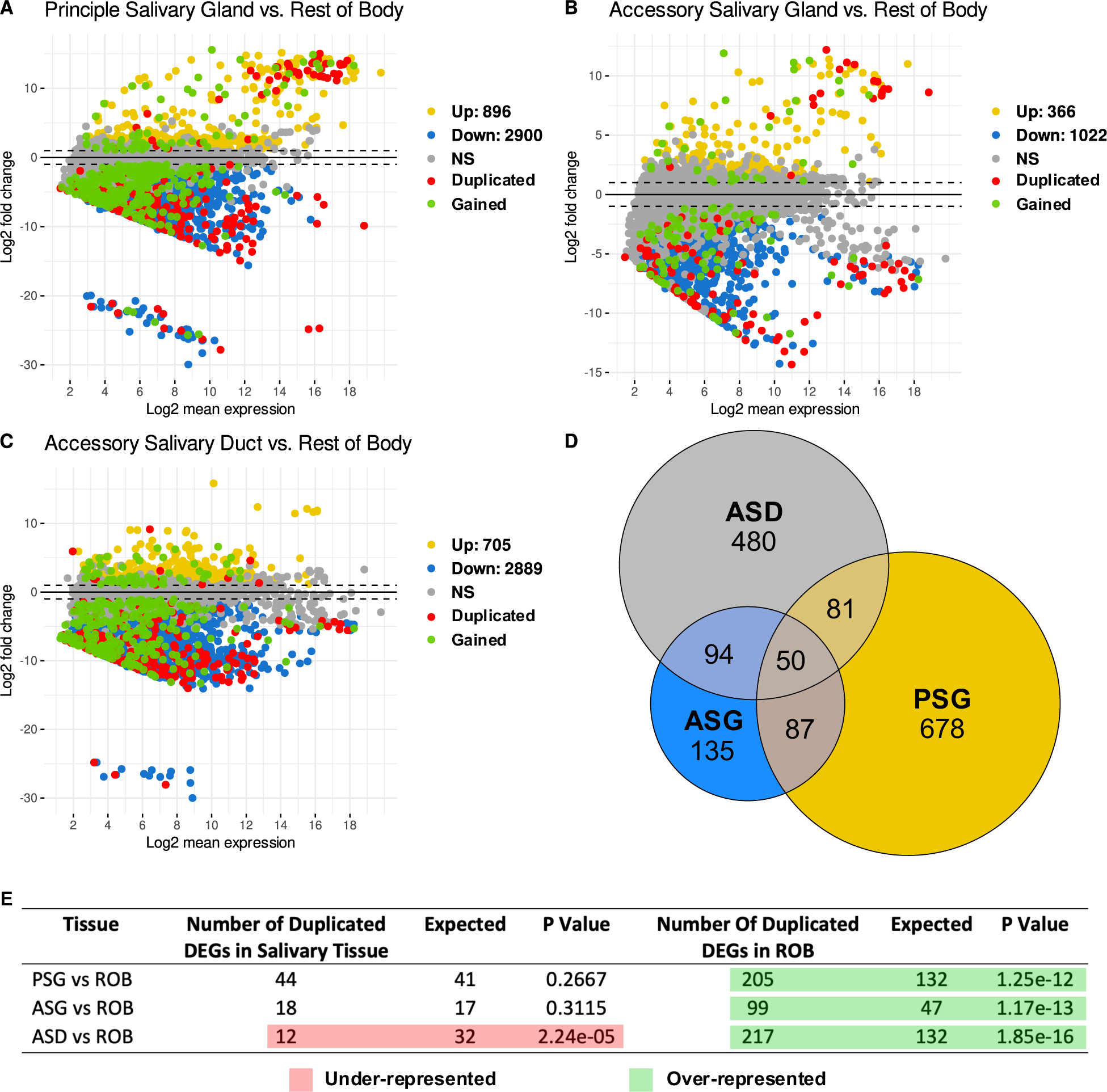
Differential expression of genes expressed in the salivary tissues versus the rest of the body. **A-C.)** MA plots for each salivary tissue vs. the rest of the body (ROB). Dotted lines indicate log2 fold change values of one and negative one. Genes upregulated in the salivary tissues are shown in yellow, and genes upregulated in the ROB are shown in blue. Points labeled in red are genes that were assigned as duplicated by pyHam analysis. Genes that are in green are genes that were assigned as gained by pyHam analysis. Genes that were not significantly differentially expressed are shown in gray. **D.)** Venn diagram showing the number of overlapping genes within each salivary tissue. The lack of shared genes suggests there are distinct roles within each salivary tissue. **E.)** Table showing the number of duplicated genes in the redbanded stink bug lineage that are also differentially expressed. Rows highlighted in red indicate that there were significantly less duplicated and differentially expressed genes than expected, while rows highlighted in green indicated that there were significantly more duplicated and differentially expressed genes than expected. Expected values were estimated using the mean of 10,000 gene overlap simulations, and statistical deviation from the expected value was measured using a hypergeometric test.

**Table 1:** GO terms significantly enriched in the genes differentially expressed in the redbanded stink bug PSG.

| GO Term ID | Gene Ontology Term | Number of Genes (Total) | P Value |
| --- | --- | --- | --- |
| GO:0003735 | structural constituent of ribosome | 58 (60) | 0.00077 |
| GO:0001216 | DNA-binding transcription activator activity | 5 (5) | 0.00263 |
| GO:0001228 | DNA-binding transcription activator activity, RNA<br>Pol II specific | 5 (5) | 0.00263 |
| GO:0008235 | metalloexopeptidase activity | 5 (5) | 0.00277 |
| GO:0008238 | exopeptidase activity | 5 (5) | 0.00277 |
| GO:0008233 | peptidase activity | 21 (22) | 0.00466 |
| GO:0008236 | serine-type peptidase activity | 6 (7) | 0.00918 |
| GO:0008237 | metallopeptidase activity | 7 (7) | 0.00918 |
| GO:0017171 | serine hydrolase activity | 6 (6) | 0.00918 |
| GO:0004252 | serine-type endopeptidase activity | 4 (5) | 0.0098 |
| GO:0019787 | ubiquitin-like protein transferase activity | 9 (9) | 0.01883 |
| GO:0140096 | catalytic activity, acting on a protein | 60 (66) | 0.02159 |
| GO:0005484 | SNAP receptor activity | 5 (5) | 0.02547 |
| GO:0019843 | rRNA binding | 20 (20) | 0.02624 |
| GO:0008270 | zinc ion binding | 13 (13) | 0.03484 |
| GO:0000049 | tRNA binding | 9 (9) | 0.03526 |
| GO:0030545 | signaling receptor regulator activity | 6 (6) | 0.03661 |
| GO:0030546 | signaling receptor activator activity | 6 (6) | 0.03661 |
| GO:0044389 | ubiquitin-like protein ligase binding | 18 (18) | 0.04635 |
| GO:0005215 | transporter activity | 36 (45) | 0.04823 |
| GO:0005198 | structural molecule activity | 69 (74) | 0.04880 |

**Table 2:**
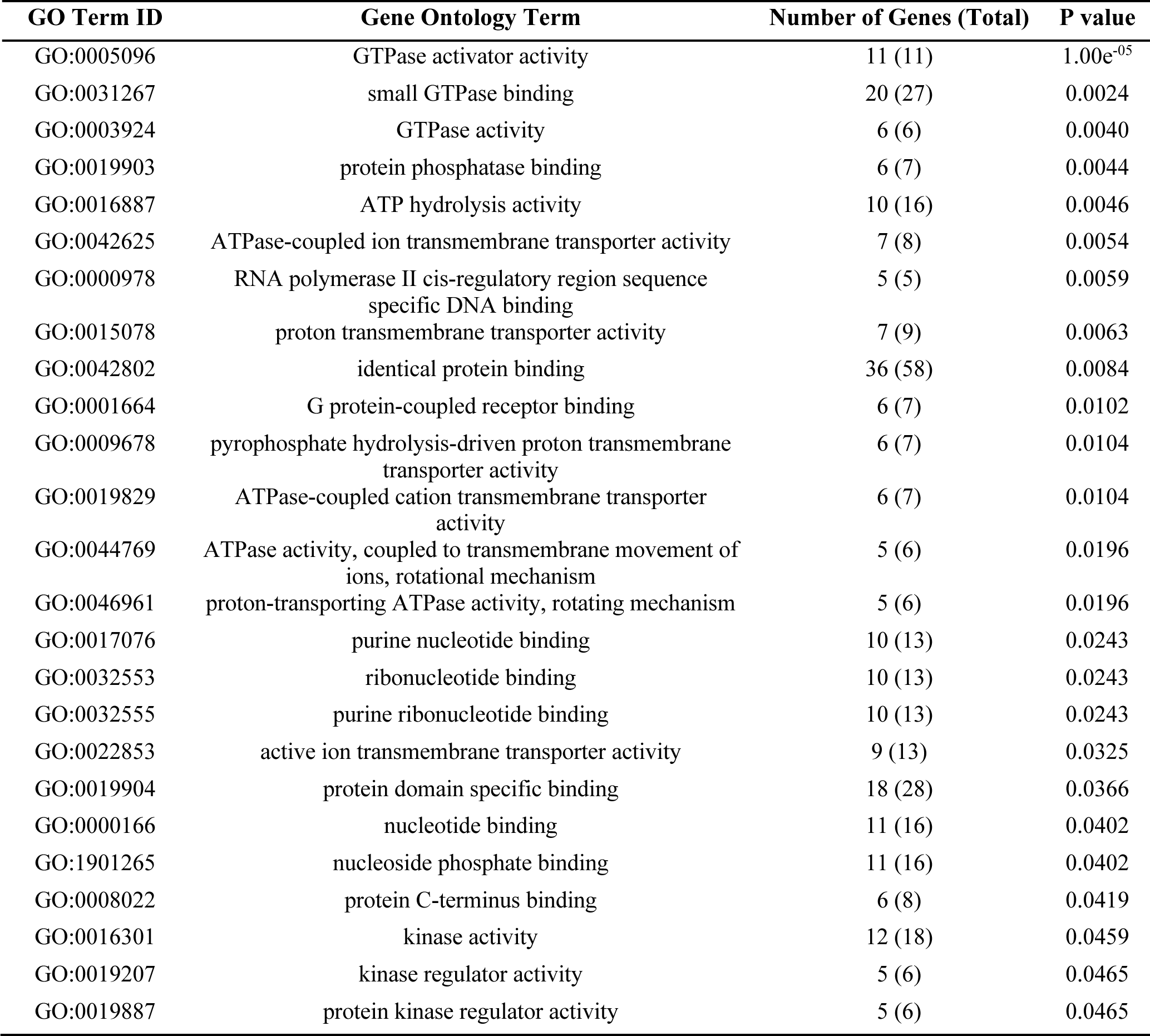
GO terms significantly enriched in the genes differentially expressed in the redbanded stink bug ASG.

**Table 3:** GO terms significantly enriched in the genes differentially expressed in the redbanded stink bug ASD.

| GO Term ID | Gene Ontology Term | Number of Genes (Total) | P value |
| --- | --- | --- | --- |
| GO:0046961 | proton-transporting ATPase activity, rotational mechanism | 8 (8) | 0.00095 |
| GO:0016301 | kinase activity | 22 (23) | 0.00194 |
| GO:0016773 | phosphotransferase activity, alcohol group as acceptor | 20 (21) | 0.00504 |
| GO:0017111 | ribonucleoside triphosphate phosphatase activity | 31 (35) | 0.01598 |
| GO:0008092 | cytoskeletal protein binding | 35 (38) | 0.01769 |
| GO:0016887 | ATP hydrolysis activity | 23 (25) | 0.02254 |
| GO:0004672 | protein kinase activity | 16 (17) | 0.03142 |
| GO:0016462 | pyrophosphatase activity | 35 (40) | 0.03230 |
| GO:0016817 | hydrolase activity, acting on acid anhydrides | 35 (40) | 0.03230 |
| GO:0016818 | hydrolase activity, acting on acid anhydrides, in phosphorus-containing anhydrides | 35 (40) | 0.03230 |
| GO:0035254 | glutamate receptor binding | 6 (6) | 0.03302 |
| GO:0016849 | phosphorus-oxygen lyase activity | 5 (5) | 0.03741 |
| GO:0050661 | NADP binding | 5 (5) | 0.04953 |

### Genes Duplicated and Differentially Expressed in the Redbanded Stink Bug

We took special interest in the genes that were duplicated and differentially expressed in the redbanded stink bug salivary gland, as we predicted that this would be the set of genes that are linked to its distinct phenotype: its higher damage potential in soybean. We found that in the ASG and PSG, there were no more duplicated and differentially expressed genes than expected at random (**Figure 4E**), and there was no significant GO term enrichment in these gene sets. In the ASD, there were fewer duplicated and differentially expressed genes than expected by chance (p = 2.24e-05) (**Figure 4E**). Similar patterns were observed in the set of genes gained and differentially expressed in the redbanded stink bug lineage, but there was functional enrichment of biological processes such as response to hormones, inorganic cation transmembrane transport, metabolic processes, and homeostatic processes in the ASD. These results suggest that gene gains or duplications are not the driver of the redbanded stink bug’s increased ability to damage to soybean relative to other stink bugs.

### Genes Retained in the Redbanded Stink Bug Lineage, but Lost in Other Pentatomids

In addition to gene gains, the differential retention of genes between lineages can have important phenotypic outcomes. To see if differential retention played a role in the redbanded stink bug’s ability to digest plant tissues, we analyzed the genes that were differentially retained in the redbanded stink bug lineage compared to the other pentatomids included in our study (*Halyomorpha halys*, *Nezara viridula*, and *Euschistus heros*). We found 2,053 genes that were lost in *H. halys* (brown marmorated stink bug), 3,486 that were lost in *N. viridula* (southern green stink bug), and 1,912 that were lost in *E. heros* (neotropical brown stink bug) but retained in the redbanded stink bug lineage. Because the PSG is the main site where extraoral digestive enzymes are synthesized, we investigated if any of the genes that were lost in other pentatomid lineages were also differentially expressed in the PSG. We found 107, 177, and 102 genes differentially expressed in the redbanded stink bug PSG that were lost in the brown marmorated stink bug, the southern green stink bug and the neotropical brown stink bug, respectively (**Figure 5**). Furthermore, we tested for GO term enrichment of biological processes, and found that all the gene sets that were lost in the other stink bug lineages and highly expressed in the redbanded stink bug were enriched with GO terms involving the breakdown of proteins and other organic molecules (**Figure 5**). This suggests that differentially retained genes in the redbanded stink bug lineage contribute to its comparatively high damage to soybean instead of genomic innovations.

**Figure 5:**
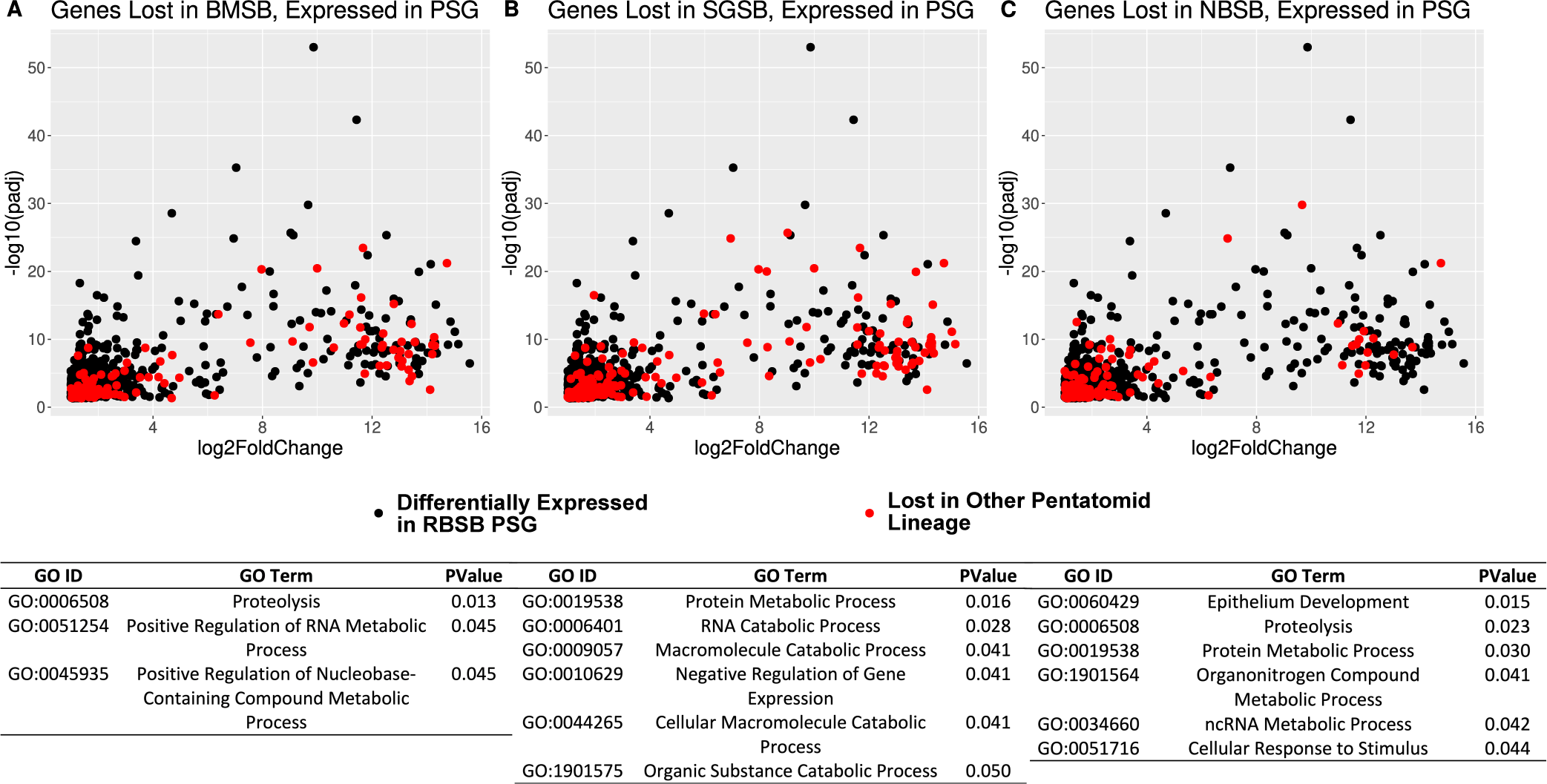
Genes that were lost in other stink bug lineages but retained and differentially expressed in the redbanded stink bug PSG. **A-C.)** Genes that were lost in the brown marmorated stink bug (BMSB), the southern green stink bug (SGSB), and the neotropical brown stink bug (NBSB), but differentially expressed in the redbanded stink bug salivary gland. Below each plot is a corresponding table of the significantly enriched biological process GO terms in the set of genes that were lost in other stink bug lineages, but differentially expressed in the redbanded stink bug PSG.

## DISCUSSION

In this study, we took a comparative genomic and transcriptomic approach to investigate the high damage potential of the redbanded stink bug with a particular interest in its distinct digestive capabilities. Previous studies have noted that the redbanded stink bug has a greater damage potential in soybean than other pentatomids due to increased chemical dissolution from salivary enzymes (Corrêa-Ferreira & De Azevedo 2002; Depieri & Panizzi 2011; Husch et al. 2014). We hypothesized that this was because the redbanded stink bug genome had undergone lineage-specific changes in its complement that led to an expanded repertoire of digestive enzymes highly expressed in the salivary gland. We found that the redbanded stink bug exhibited similar gene turnover rates to the rest of the Pentatomidae (λ_rbsb_ = 0.0022, λ_p_ = 0.0026), suggesting that the redbanded stink bug genome is not evolving more rapidly than the other pentatomids included in our study. However, we did find significant enrichment of genes with digestive capabilities such as trypsins, ovochymases, and cathepsins in the set of genes duplicated in the redbanded stink bug (**Figure 3**), and we also found high expression of genes involved in these processes in the salivary tissues (**Figure 4**, **Tables 1-3**).

Considering this, we identified the set of genes that were duplicated and differentially expressed in the salivary gland. We found that there was no difference in the number of duplicated and differentially expressed genes we observed and the number expected by chance in the PSG and ASG, and significantly less than expected in the ASD (**Figure 4E**). Furthermore, we did not find any significant functional enrichment in the set of duplicated genes that were also differentially expressed in the salivary glands. By contrast, we did find significantly more duplicated genes differentially expressed in the ROB than expected (**Figure 4E**). In the set of duplicated and differentially expressed genes in the ASD vs. ROB, we found significant functional enrichment for multiple types of peptidase activity in the ROB. This is likely because the gut, another major site of digestion, was kept in the ROB tissues. Furthermore, the peptidase profiles were different between the salivary tissues and the ROB, as no cysteine-type endopeptidases were detected in the salivary tissues, but many were detected in the ROB. This is consistent with previous studies on the southern green stink bug salivary and gut proteases (Lomate & Bonning 2016). Previous work has suggested that some gut digestive enzymes can be transported to the saliva and may also play a role in extraoral plant tissue digestion, but our study did not account for this (Liu & Bonning 2019).

Because we did not find substantial evidence of increased gene dosage of digestive genes expressed in the redbanded stink bug salivary gland, we also considered the genes that were differentially retained in the redbanded stink bug lineage. Interestingly, we found enrichment of functions involved in proteolysis and the breakdown of organic molecules in the genes that are differentially retained and expressed in the stink bug PSG (**Figure 5**). In a study comparing soybean seed damage of four species of stink bugs, Depieri and Panizzi (2011) observed that the redbanded stink bug inflicted significantly more damage to soybean seeds than the other species (including *N. viridula* and *E. heros*), to the point where the protein bodies within the seeds were completely dissolved. Our results suggest that this extreme dissolution of the soybean protein bodies could be attributed to the differential retention of genes instead of gene gain or duplication in the redbanded stink bug lineage.

Along with serious damage to plant tissues, redbanded stink bugs are known to be resilient to pesticide application (Baur et al. 2010; Temple et al. 2013b). We found an overrepresentation of genes involved in detoxification processes that were duplicated in redbanded stink bug lineage including cytochrome P450s, carboxylesterases, glutathione S-transferases, and ATP-binding cassette (ABC) transporters in the set of duplicated genes in the redbanded stink bug genome (**Figure 2A**). Many of these genes were also differentially expressed in the ROB, which is expected as detoxification is widely carried out in tissues other than the salivary glands (Heidel-Fischer & Vogel 2015).

We found a set of genes that were lost in other stink bug lineages but retained and differentially expressed in the redbanded stink bug salivary gland, but we did not find strong support for the duplicated or gained genes being the enzymes responsible for the redbanded stink bug’s high digestive capabilities. However, the differentially retained salivary genes are enriched with protein families involved in proteolysis, and thus could be the enzymes responsible for the redbanded stink bug’s high damage. This could be related to the redbanded stink bug’s more narrow host range than other stink bug pests, as it has been noted that the redbanded stink bug mainly feeds on legumes, while other stink bugs will feed on other crops such as cotton, corn, and rice (Temple et al. 2013a). We speculate that the genes lost in the other stink bug lineages were not beneficial for a generalist diet and may have eroded from the genomes of these lineages, while they were retained in the redbanded stink bug lineage as it continued to feed on protein-rich legumes. Our work identifies several candidate genes for lab studies and proposes an interesting link between genomic history and the redbanded stink bug’s agriculturally relevant physiology.

## MATERIALS AND METHODS

### Collection, Dissection, and RNA extraction

Redbanded stink bug samples were collected from an experimental soybean field at the Macon Ridge Research Station in Winnsboro, Louisiana, between May 2021 and September 2022. They were collected using 38.1 cm diameter sweep nets (BioQuip). We transported them to a 30.48 × 30.48 × 30.48 cm BugDorm (BioQuip) within a growth chamber (Percival, Iowa, USA) at Mississippi State University. The rearing chamber was maintained at 28° C and 80% relative humidity with a 12:12 (L:D) photoperiod and the redbanded stink bugs were fed a diet of organic green beans. We dissected adult stink bugs in cold phosphate buffered saline (PBS) under a stereomicroscope (Zeiss SteREO Discovery, Oberkochen, Germany). We separated them into four tissues: the principal salivary glands (PSG), accessory salivary glands (ASG), accessory salivary duct (ASD), and the rest of the body without salivary tissue (ROB) (**Figure 1**) and immediately placed them in RNAzol (Molecular Research Center, Cincinnati, OH, USA) in pools of three individuals. We made four replicates of each tissue sample, further separating two male replicates and two female replicates, although we do not expect large differences in salivary gene expression between male and female. We isolated total RNA using standard RNAzol protocol, and further purified samples using a NEB Monarch^®^ RNA Cleanup kit (New England Biolabs, Ipswich, MA, USA). We checked total RNA quantity and quality using a NanoDrop One spectrophotometer (Thermo Fisher Scientific, Waltham, MA, USA) and agarose gel electrophoresis. We also assessed RNA integrity using an Agilent 2100 Bioanalyzer (Agilent, Santa Clara, CA, USA).

### Library Preparation and Sequencing

We prepared sixteen sequencing libraries using standard Illumina library prep with poly-A enrichment for mRNA and sequenced them using a NovaSeq 6000 instrument generating 150 bp paired-end reads. We checked read quality using FastQC v0.11.9 (Andrews 2010) and trimmed adapters and low quality reads with Trimmomatic v0.39 (Bolger et al. 2014). The remaining reads were mapped to the SILVA rRNA database (Quast et al. 2013) to remove any ribosomal RNA contamination using Bowtie2 v2.4.4 (Langmead & Salzberg 2012).

### Genome Annotation

We masked the repetitive elements of the redbanded stink bug genome (GCA_023052935.1) using RepeatModeler v2.0.3 and RepeatMasker v4.1.3 (Tarailo-Graovac & Chen 2009; Flynn et al. 2020), and mapped the trimmed RNA-seq reads to the masked genome using HISAT2 v2.2.1 (Kim et al. 2019). We converted the output to a sorted BAM file using Samtools v1.6 (Li et al. 2009) and used the resulting alignment file to annotate the genome with the BRAKER2 v2.1.6 pipeline (Brůna et al. 2021). BRAKER2 allows for protein datasets from closely related species to be used for genome annotation, so we downloaded the arthropod protein dataset from OrthoDB (Waterhouse et al. 2013) and annotated the genome separately with protein data. We used TSEBRA as a transcript selector for both the protein-mapped dataset and the RNA-seq dataset (Gabriel et al. 2021). We checked the completeness of all predicted proteomes (before and after TSEBRA selection) using BUSCO v5.4 (Simão et al. 2015) in protein mode against the hemiptera_odb10 database. Finally, we filtered the annotation with the highest BUSCO score using the supplementary script provided by BRAKER selectSupportedSubsets.py using the –anysupport flag to only retain the transcripts that had external support during the annotation. We functionally annotated the genome using the resulting proteome and diamond (v2.0.15) BLASTp against NCBI’s nr protein database (https://blast.ncbi.nlm.nih.gov/Blast.cgi) and eggNOG v2.1.9 in diamond mode (Cantalapiedra et al. 2021; Buchfink et al. 2015, 2021).

### Comparative Genomics Analysis

We downloaded various hemipteran proteomes from multiple sources, listed in **Supplementary Table 1**. Each protein was represented by its longest isoform. We checked each proteome for completeness using BUSCO analysis, and only used those that were greater than or equal to 85% complete and having less than 10% duplicated single-copy orthologs. For our sampling strategy, we aimed to have a high taxonomic representation within the Pentatomorpha, as the family that stink bugs belong to, Pentatomidae, lies within this suborder. Outside of the Pentatomorpha, we used the two most divergent taxa (when available) as representatives within each infraorder or suborder. The final taxa used for comparative genomics analysis included *Acyrthosiphon pisum*, *Bemisia tabaci*, *Nilaparvata lugens*, *Apolygus lucorum*, *Rhodnius prolixus*, *Oncopeltus fasciatus*, *Riptortus pedestris*, *Halyomorpha halys*, *Nezara viridula*, *Euschistus heros*, and *Piezodorus guildinii.* For species tree construction and assessment of comparative genomics statistics, we used OrthoFinder v2.5.5 using the –msa flag (Emms & Kelly 2019). To assign genes to hierarchical orthologous groups (HOGs) and assess evolutionary history of gene families (i.e., gene duplications, gene gains, gene losses, etc.) we used OMA standalone v.2.5.0 and pyHam, respectively (Altenhoff et al. 2019; Train et al. 2019). To test for rapid expansions and contractions in the HOGs inferred from OMA, we used CAFE5 (Mendes et al. 2020). We made the species tree ultrametric using the supplemental orthofinder script make_ultrametric.py, calibrating the root of the tree at 386 million years ago (MYA) (Johnson et al. 2018). To ensure optimal parameters, we ran CAFE5 employing error model estimation and iteratively increased the number of gamma rate categories (-k) until negative log-likelihood stopped improving. To test if the redbanded stink bug has a higher rate of gene turnover compared to the other taxa in our study, we used CAFE5 to estimate separate gene turnover rates (λ) for the redbanded stink bug lineage (λ_rbsb_), the Pentatomidae (λ_p_) and the rest of the Hemiptera (λ_h_).

To determine functional enrichment of the genes that were duplicated, gained, and lost, we conducted enrichment analysis with clusterProfilr v.4.6.2 using KEGG orthology terms annotated with eggNOG (Wu et al. 2021; Kanehisa et al. 2016). To retrieve the genes that were duplicated, gained, and lost in the redbanded stink bug lineage, we conducted a vertical genome comparison between the extant redbanded stink bug genome and its most recent ancestor and used pyHam’s .get_duplicated(), .get_gained, and .get_lost(), functions. To retrieve the genes that were lost in other pentatomid lineages but retained in the redbanded stink bug lineage, we conducted horizontal genome comparisons between the redbanded stink bug and the other lineages separately, and used the .get_lost() function to retrieve the gene names. To test for functional enrichment in the genes that were duplicated, gained, and retained in redbanded stink bug lineage, we used the annotated redbanded stink bug genome as the set of background genes. For genes that were lost in the redbanded stink bug lineage, we used pyHam’s get_ancestral_genome_by_name() function to reconstruct the genome of the most recent common ancestor of the redbanded stink bug, the brown marmorated stink bug, and the southern green stink bug and completed the genome by selecting one representative gene from each HOG returned by the ancestral genome’s .genes list. We annotated the ancestral genome using eggNOG and conducted enrichment analysis using the ancestral genome as the background set of genes. We matched the genes from lost HOGs in the redbanded stink bug genome to the reconstructed ancestral genome and used them for enrichment analysis in clusterProfilr.

### Differential Expression Analysis

We mapped each read dataset to the annotated genome using STAR v2.7.10b (using the Quantmode option) (Dobin et al. 2013) and the subset of supported annotations from the BRAKER2 pipeline. We measured transcript abundance using RSEM v1.3.3 (Li & Dewey 2011). We imported transcript abundance to R using tximport v1.26.1 (Soneson et al. 2015) and conducted differential gene expression analysis using DEseq2 v1.38.3, filtering out genes with less than 10 counts in greater than or equal to 4 samples (Love et al. 2014). We conducted gene ontology (GO) analysis on the significantly upregulated and downregulated genes in each tissue. Only the expressed genes (i.e., the genes that were kept for differential expression analysis) were used for the set of background genes, and GO annotations were used from the previous functional annotation of the genome with eggNOG. We tested for significant enrichment of GO terms using topGO v2.50.0 (Alexa & Rahnenfuhrer 2020), employing the Kolmogorov-Smirnov test with the “elim” algorithm. To test if there were more (or less) duplicated genes that were also differentially expressed than would be expected by chance, we used a hypergeometric test implemented in the R using the “phyper” function (R Core Team 2020).

### Construction of *Piezodorus* Phylogeny

To investigate phylogenetic relationships within the genus *Piezodorus*, we downloaded 58 *Piezodorus cytochrome c oxidase subunit I* (COI) sequences from the BOLD systems database (Ratnasingham & Hebert 2007), along with one *Halyomorpha halys* sequence for the outgroup. We aligned the nucleotide sequences using MAFFT v7.490 and discarded low quality and redundant sequences (Katoh & Standley 2013). We found the best nucleotide substitution model using the modelfinder algorithm incorporated in IQ-TREE v2.0.7 (Minh et al. 2020; Nguyen et al. 2015). We built a maximum-likelihood tree using the alignment implementing the best substitution model, and branch support was measured using ultrafast bootstrap with 1000 replicates, the Shimodaira-Hasegawa-like approximate likelihood ratio test (SH-aLRT) with 1000 replicates, and the aBayes test (Anisimova et al. 2011; Minh et al. 2013; Hoang et al. 2018)

## FUNDING

This work was supported by a Foundation for Food and Agriculture Research New Innovator Award [534275 to J.G.K].

## ACKNOWLEDGEMENTS

The authors would like to thank Matthew J. Ballinger, Ashli E. Brown, Amy L. Dapper, and Andy D. Perkins for their helpful comments on the manuscript. We also thank Travis C. van Warmerdam for taking the photograph depicted in **Figure 1A**.

## AUTHOR CONTRIBUTIONS

Conceptualization: H.K.W. and J.G.K.; Methodology: H.K.W.; Formal Analysis: H.K.W.; Investigation: H.K.W., J.G.K., and F.G.H.; Resources: J.G.K., T.B.T, and F.G.H.; Writing – Original Draft: H.K.W.; Writing Review & Editing: H.K.W., J.G.K., T.B.T., S.A., and F.G.H.; Funding Acquisition: J.G.K.; Supervision: J.G.K. and F.G.H.

## SUPPLEMENTARY MATERIAL

Supplementary data will be available online.

## DATA AVAILABILITY STATEMENT

All reads generated for this study were deposited to NCBI’s sequence read archive under BioProject accession PRJNA729517. Accession numbers or links to all genomes used in this study are provided in **Supplementary Table 1.**

## Notes

### Competing Interest Statement

The authors have declared no competing interest.

