## Supplemental table 1 and supplemental figures 1-2 for "Comparative genomics and the salivary transcriptome of the redbanded stink bug shed light on its high damage potential to soybean"

**SUPPLEMENTARY DATA**

**Supplementary Table 1: Summary of genomes screened for use in our comparative genomics analysis.** The genomes in bold text did not pass our filters (≥85% BUSCO completeness, <10% BUSCOs duplicated)

| Species Name | Database & Accession | BUSCO proteome completeness (%) | BUSCO proteome duplicated (%) |
|---|---|---|---|
| ***Aspongopus chinensis*** | **https://figshare.com/s/7bc3ec0b7fff879ecf6e** | **58.7** | **1.8** |
| *Acyrthosiphon pisum* | Refseq: GCF_005508785.2 | 98.8 | 2.5 |
| *Apolygus lucorum* | Genbank: GCA_009739505.2 | 93.9 | 5.4 |
| *Bemisia tabaci* | Refseq: GCF_001854935.1 | 98.8 | 1.7 |
| *Euschistus heros* | Genbank: GCA_003667255.2 | 94.9 | 3.9 |
| *Halyomorpha halys* | Refseq: GCF_000696795.2 | 99.2 | 1.9 |
| ***Homalodisca vitripennis*** | **Refseq: GCF_021130785.1** | **92.8** | **10.8** |
| *Nezara viridula* | Genbank: GCA_928085145.1 | 90.3 | 2.0 |
| *Nilparvata lugens* | Refseq: GCF_014356525.2 | 97.0 | 5.6 |
| *Oncopeltus fasciatus* | https://i5k.nal.usda.gov/Oncopeltus_fasciatus | 88.3 | 1.6 |
| *Piezodorus guildinii* | Genbank: GCA_023052935.1 | 85.4 | 1.0 |
| *Riptortus pedestris* | https://doi.org/10.5281/zenodo.4766420 | 97.3 | 2.2 |
| *Rhodnius prolixus* | https://vectorbase.org/vectorbase/app/record/dataset/DS_b8c0427e28 | 91.8 | 1.3 |

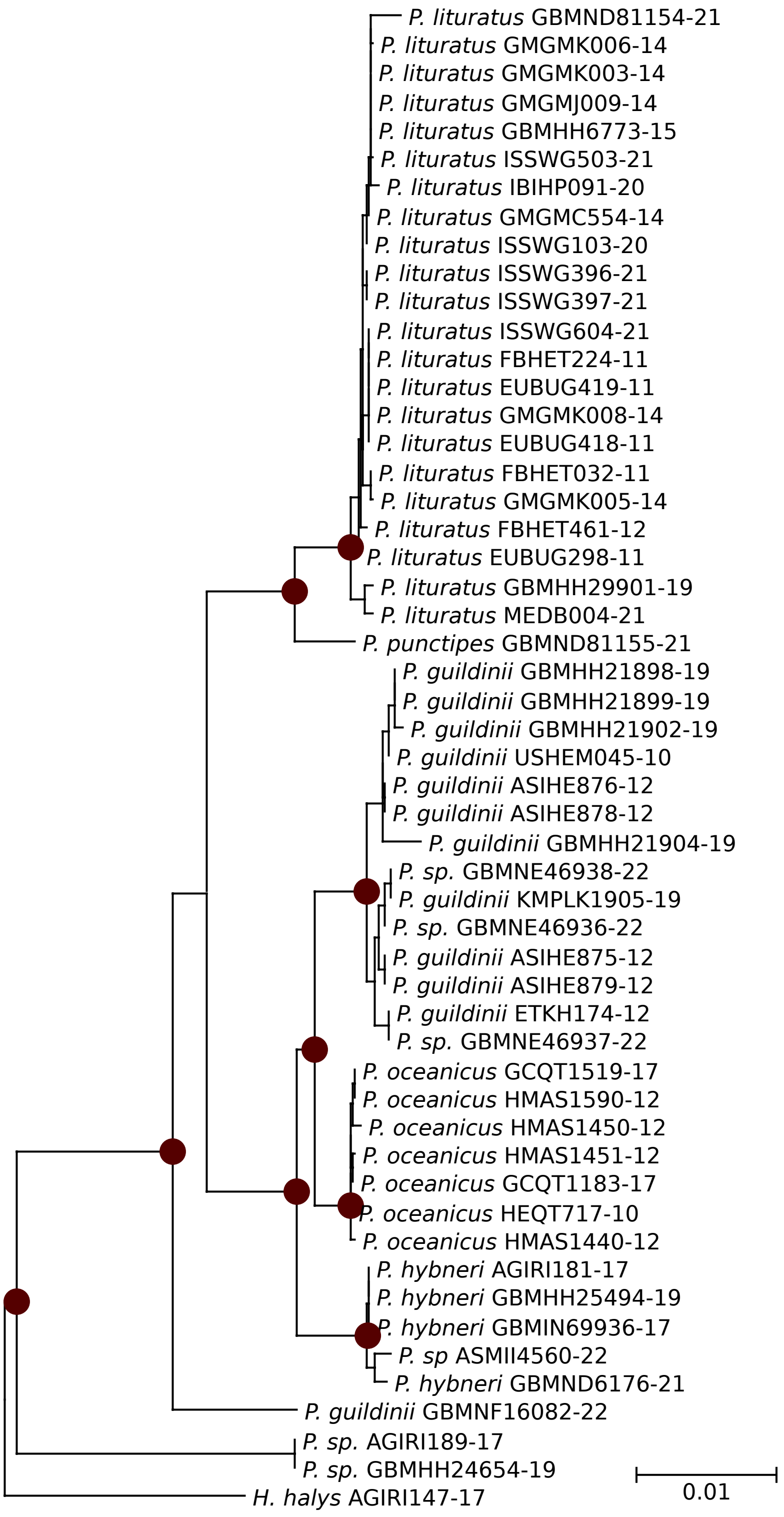


**Supplementary Figure 1: Phylogeny of genus *Piezodorus* based on *cytochrome b oxidase subunit I* (COI).** Maroon dots indicate bootstrap support ≥ 75 at relevant nodes.

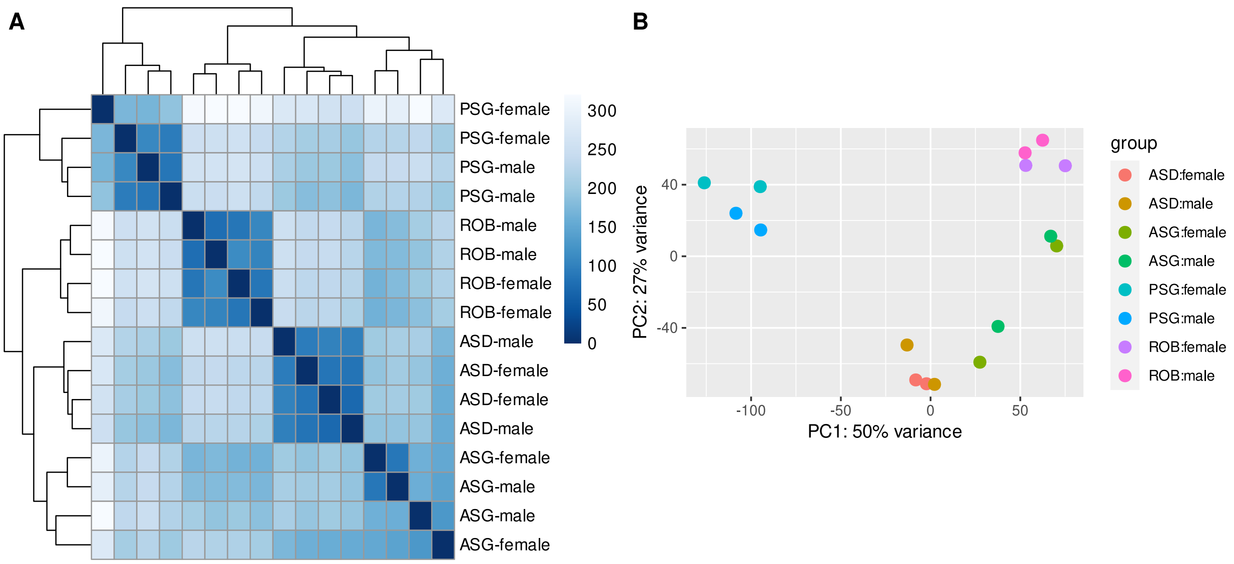


**Supplementary Figure 2: Correlations of differential expression between datasets. A.)** Heatmap and hierarchical clustering of distances between differential expression datasets separated by tissue and sex. **B.)** Principal component analysis of differential expression datasets. The datasets cluster by tissue, but not by sex.
